# The C-terminal domain of transcription factor RfaH: Folding, fold switching and energy landscape

**DOI:** 10.1101/2020.09.26.315077

**Authors:** Bahman Seifi, Stefan Wallin

## Abstract

We study the folding and fold switching of the C-terminal domain (CTD) of the transcription factor RfaH using a hybrid sequence-structure based model. We show that this model captures the essential thermodynamic behavior of this metamorphic domain, i.e., a switch in the global free energy minimum from an *α*-helical hairpin to a 5-stranded *β*-barrel upon separating the CTD from the rest of the protein. Using this model and Monte Carlo sampling techniques, we analyze the energy landscape of the CTD in terms of progress variables for folding towards the two folds. We find that, below the folding temperature, the energy landscape is characterized by a single, dominant funnel to the native *β*-barrel structure. The absence of a deep funnel to the *α*-helical hairpin state reflects a negligible population of this fold for the isolated CTD. We observe, however, a significantly higher *α*-helix structure content in the unfolded state compared to results from a similar but fold switch-incompetent version of our model. Moreover, in folding simulations started from an extended chain conformation we find transient *α*-helix structure that disappears as the chain progresses to the thermally stable *β*-barrel state.

## 1 Introduction

Proteins are increasingly being discovered with a remarkable ability to switch from one native fold into another one with widely different structure [1–3]. While it is not uncommon for proteins to undergo large-scale motions after their initial folding, such as domain-swapping [4] or other hinge-like motions [5], fold switching is a distinct phenomenon. By definition, it involves a reorganization of the protein at the most basic structural level at play in folding, i.e., secondary structure (*α*-helices and *β*-sheets). Despite the remarkable complexity of these molecular transformations, fold switching is reversible and thereby controlled by the system’s free energy. In this sense, metamorphic proteins can be said to adhere to Anfinsen’s thermodynamic principle (or hypothesis) of protein folding [6]. Clearly, however, fold switching fundamentally challenges the idea of a “unique” native conformation, which was a central aspect of the classic view of folding since emerging from the pioneering refolding experiments on ribonuclease A [7]. It is important to note that fold switching typically occurs only when triggered by specific changes to the local environment or milieu of the protein, such as salt concentration [8], redox condition [9] or oligomerization state [10]. In the absence of such a trigger, metamorphic proteins typically fold to an apparently unique structure, which masks their fold switching capabilities. As a result, metamorphic proteins often go unrecognized [11].

Metamorphic proteins thus encode two different folds within a single amino acid sequence even though, as mentioned, they typically adopt a single fold for a given (constant) local milieu. It is natural, then, to ask: what impact does this dual encoding have on their folding? This question is in fact related to a classic line of inquiry in protein folding, namely whether the mechanism of folding is conserved among homologous proteins or, more generally, among sequences adopting the same fold [12, 13]. It was observed, remarkably, that sequences with low sequence similarity (but still adopting the same native fold) often fold in a very similar manner [14–16], however, this conservation break down at very low sequence similarity [17] and does not extend to all fold classes [17–19]. The sequences of metamorphic proteins have diverged from their homologs to the point of nearly adopting a new unique fold. Characterizing the folding of metamorphic proteins may shed light on the limits of how the amino acid sequence can shape the energy landscape of proteins. It may also be practically important in the field of protein metamorphism because any identifying feature, including folding behavior, can be used to discover as of yet unknown fold switching events [10, 11, 20].

Here we study the C-terminal domain (CTD) of the transcriptional antiterminator protein RfaH, a prototypical metamorphic protein [21]. This fold switching exhibited by this protein has been studied extensively both experimentally [22–25] and computationally [26–33]. The RfaH CTD, on its own, i.e., excised from the rest of protein, folds spontaneously into a *β*-barrel-like fold with 5 strands, as shown in Fig. 1. This structure is virtually identical to the CTD structure of other members of the NusG/Ste5 family of transcription factors to which RfaH belongs [34]. However, as part of the full-length RfaH, the CTD adopts instead an *α*-helical hairpin fold [22]. This entirely different folded state is stabilized by favorable interactions with the RfaH N-terminal domain (NTD) [23]. The switch in structure from all-*α* to all-*β* is triggered when RfaH binds to RNA polymerase in a paused state, which underpins RfaH’s regulatory function in transcription and translation [24, 35].

**Figure 1:**
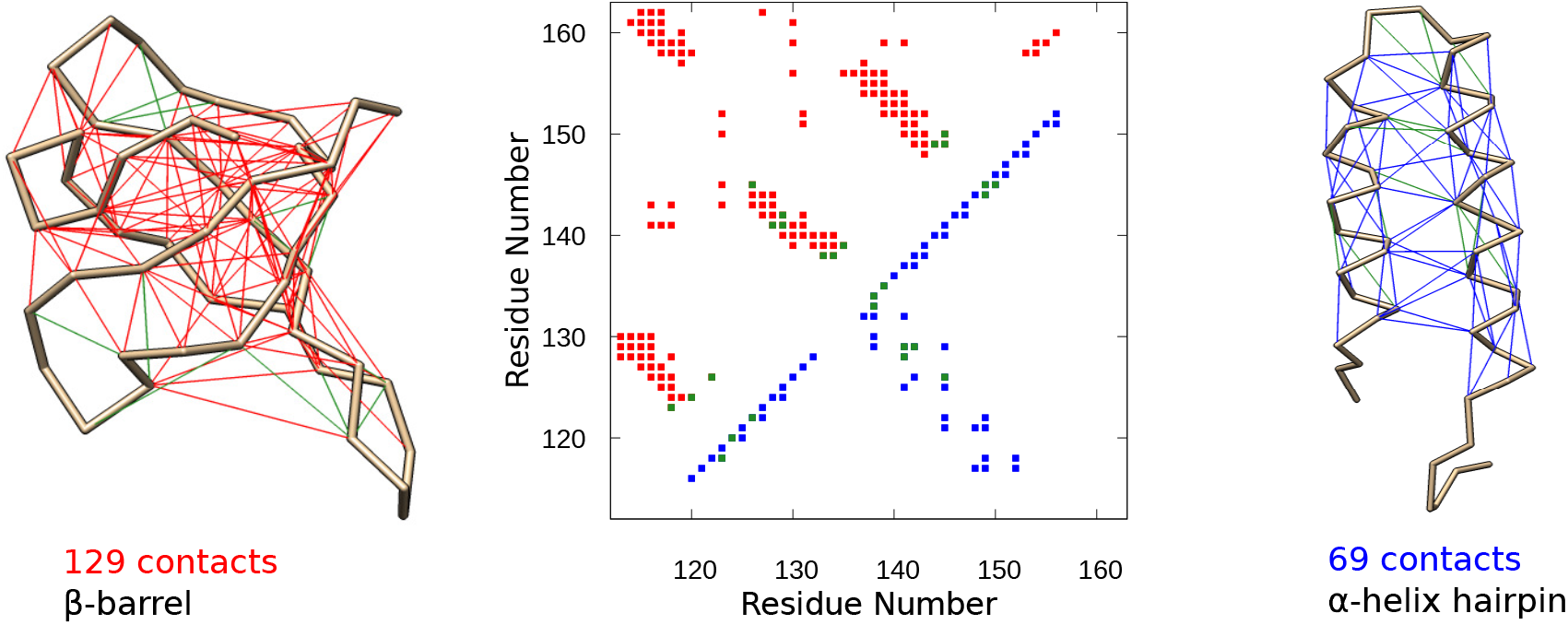
The two different folds of the C-terminal domain (CTD) of RfaH. Native contacts in the two folds, determined using the shadow map method [54], are shown as thin lines in the C_*α*_-traces (left and right) and as solid squares in the contact maps (center). The two contact sets, denoted C^*β*^ (*β*-barrel) and C^*α*^ (*α*-helical hairpin), have 13 common contacts (green).

Simulating the structural transitions of metamorphic proteins on the computer has the potential for detailed insight but is technically challenging. Some studies have focused on alleviating the formidable problem of achieving sufficient conformational sampling for these large-scale transitions by various enhanced sampling techniques [26, 28, 32, 36, 37]. This approach assumes, however, an underlying model that accurately captures the delicate free energy balance between folds. This is not guaranteed even for modern explicit-water molecular dynamics force fields [38], although it has been achieved within the framework of coarse-grained 3-letter protein model [39–41]. Other studies have relied on so-called structure-based models (SBM). The basic idea of SBMs is to make the contacts present in a given target (typically native) structure artificially favorable, while non-native interactions are left neutral or even made repulsive [42]. SBMs can be constructed with several basin of attractions, each one representing a different target structure. Single-basin SBMs have a long tradition in protein folding studies [43] and has had notable successes in reproducing detailed experimental data on specific proteins, such as folding rates [44, 45] and transition-state structures [46], despite the *ad hoc* nature of the approach. Dual- or multi-basin SBMs are the logical extension to metamorphic proteins, and such models have been applied to both the CTD of RfaH [27, 30] and the mutation-induced fold switch between the GA and GB proteins [47, 48].

To study the energy landscape of the RfaH CTD, we take a hybrid approach that combines a physics-based model for protein folding with a dual-basin SBM. We aim to develop this hybrid sequence-structure based model based on its basic thermodynamic behavior, which we determine using extensive Monte Carlo simulations. Hence, we require that this model captures the switch in global free energy minimum between the two folds of RfaH CTD, depending on whether it is part of the full-length RfaH or considered as an isolated fragment. We also consider a single-basin version of our model that does not exhibit proper fold switching. By comparing closely with this model, we probe the impact of encoding for two different folds on the energy landscape of RfaH CTD.

## 2 Materials and Methods

### 2.1 Physics-based computational protein model

All simulations are carried out using the software package PROFASI [49]. The physics-based model implemented in this package is described in Ref. [50]. Briefly, the model combines an all-atom protein representation with an effective potential energy function with 4 terms: *E*^(0)^ = *E*_loc_ + *E*_ev_ + *E*_hb_ + *E*_sc_. Solvent molecules are implicitly taken into account by the energy function. The term *E*_ev_ implements excluded-volume effects between all atom pairs using a 1/*r*^12^-function. The term *E*_loc_ includes interactions between atoms close along the chain, e.g., between partial charges on neighboring peptide planes, which help provide a good local chain description. The remaining two terms, *E*_hb_ and *E*_sc_, represent hydrogen bonding and sidechain-sidechain interactions, respectively. Hydrogen bonding is implemented through orientationally dependent attractions between donor and acceptor groups. The term *E*_sc_ includes both pairwise interactions between partial charges on sidechains and effective hydrophobic attractions.

### 2.2 Monte Carlo simulations

Simulated tempering Monte Carlo simulation were used to determined equilibrium behavior (thermodynamics). Both global (pivot) and small-step (Biased Gaussian Steps [51] and sidechain rotations) Monte Carlo moves are included to speed up efficient sampling of conformational space. Our small-step “kinetic” Monte Carlo runs differ from our simulated tempering runs in two ways: (1) global MC moves are turned off and (2) the temperature is held fix. Statistical errors are estimated from independent simulations. Thermodynamic simulations of the isolated RfaH CTD are obtained using 20 independent runs of each 3 × 10^7^ MC cycles, where each cycle is 239 elementary MC moves (the number of turnable torsion angle of the simulated chain). For the full-length RfaH, 10 independent runs of at least 1 × 10^7^ MC cycles, where a cycle is 740 elementary steps.

### 2.3 Representative experimental structures

As representative structures of the two RfaH folds, we use the X-ray structure of the fulllength RfaH with PDB id 2oug [22] and the NMR derived structure of the isolated CTD with PDB id 2lcl [21]. The missing residues in 2oug, including the linker region (101-114) and flexible C-terminal tail (157-162), were added back using a homology modeling tool [52], as described previously [33]. The 2lcl structure of the isolated CTD was truncated to retain the ordered part (residues 113-162).

### 2.4 Native contact maps

The representative structures were submitted to the SMOG webserver [53] to obtain sets of residue-residue contacts according to the shadow map algorithm [54]. For 2lcl, we obtained a set of 129 native contacts, C^*β*^. For 2oug, retaining only contacts with both i and j within the segment 115-156, we obtained a set of 69 contacts, C^*α*^. The C^*α*^ and C^*β*^ contact maps are shown in Fig. 1 and used for the simulations of the isolated CTD. Simulations of the full-length RfaH are carried out with the contact map C^*α*_RfaH_^, with 157 contacts. C^*α*_RfaH_^ includes all contacts in C^*α*^ and, in addition, all the NTD-CTD inter-domain contacts, i.e., contacts between one residue in the segment 1-100 and another in the segment 115-156.

### 2.5 Observables

The progress variables, *Q_α_* and *Q_β_*, are the fraction of the native contacts formed in C^*α*^ and C^*β*^, respectively (see Fig. 1). A contact between residues i and j is considered formed if 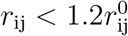, where *r*_ij_ is the distance between residues i and j (as defined in section 3.1), and 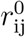 is the value rjj in the representative structure. The root-mean-square deviations, RMSD_*α*_ and RMSD_*β*_, are taken with respect to two representative structures of the all-*α* and all-*β* CTD folds, respectively, and determined over their C_*α*_ atoms.

### 2.6 Correction term for the dual-basin SBM

In our dual-basin SBM, the idea is to include attractions between pairs of residues that contact each other in the all-*α* fold (C^*α*^) and attractions between pairs of residues that contact each other in the all-*β* fold (C^*β*^). However, C^*α*^ and C^*β*^ have 13 common contacts, as shown in Fig. 1, and it would be unreasonable to include both attractions for these contacts. Therefore, we use the convention that, for each of these 13 contacts, only the most favorable of the two contact energies contributes to the total energy. This is achieved by a correction term included in the potential energy function of the dual-basin SBM (see section 3.3), which is given by

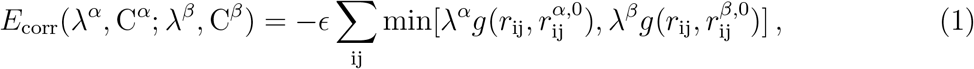

where the sum goes over the 13 contacts ij that are present in both C^*α*^ and C^*β*^. In this equation, the distance *r*_ij_, the function *g* and the strengths λ^*α*^ and λ^*β*^ are defined as in sections 3.2 and 3.3. The “native” distances 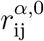 and 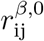 are the values of *r*_ij_ in the all-*α* and all-*β* representative structures, respectively.

## 3 Results and Discussion

### 3.1 Hybrid sequence-structure-based simulation approach

An ideal computational model for simulating metamorphic proteins would be capable of capturing the large-scale structural transitions of these proteins and, at the same time, be computationally tractable. However, as mentioned in Introduction, such an approach is out of reach at present. We pursue here a hybrid approach in which an implicit solvent physics-based model is augmented with an SBM. Similar approaches have been tested before [55–57], although not for the system under study here. Specifically, we construct the potential energy function of the hybrid model as a liner combination of the physics-based and the structurebased potentials. Our aim is to pick the relative strength of the SBM as small as possible, while requiring that the hybrid model as a whole exhibits a thermodynamic behavior in agreement with available experimental data.

As our starting point, we use the protein model developed in Ref. [50]. This model is based on an effective all-atom (solvent free) potential energy function, which includes terms for the major driving forces of protein structure formation, including hydrophobic and electrostatic attractions and hydrogen bonding. Parametrization of this model was done by requiring that a set of 17 different amino acid sequences exhibit global free energy minima corresponding to their respective experimentally determined native structures. Interestingly, this “top-down” approach to parameterization also lead to thermodynamic behaviors, such as melting temperatures, that for many of the sequences were in quantitative agreement with experimental data [50].

With this model in mind as a baseline, we formulate a structure-based, or Gō-like [43], potential, which provides an energetic bias towards a single target structure defined by a set of residue-residue contacts, C (the contact map of the structure). We pick this energy term to have the form

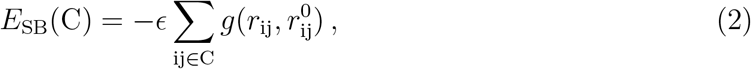

where the sum goes over all contacts ij in C, *ϵ* is the energy unit of our baseline model [50], *r*_ij_ is the C_β_-C_*β*_ distance between residues i and j, and 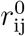 is the value of *r*_ij_ in the native fold. For any contact ij for which one (or both) positions is a glycine residue, thus lacking a C_*β*_ atom, *r*_ij_ is instead instead determined using the C_*α*_-atom position. The quantity 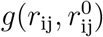, where

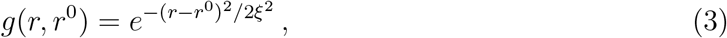

measures the extent to which the native contact ij is correctly formed. The width of the Gaussian function *g* is controlled by the parameter *ξ*, which we set to *ξ* =1 Å. Previous work [55–57] that have similarly combined physics-based and SBMs, have exclusively formulated their SBMs using C_*α*_-C_*α*_ distances. Here we primarily use C_*β*_-atoms for quantifying native contact formation because we expect that C_*β*_-C_*β*_ distances, through the function 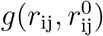, to provide a higher specificity towards the native structure.

In the following, we apply Eq. 2 to the two possible folds of the RfaH CTD. This gives two structure-based potentials, *E*_SB_(C^*α*^) and *E*_SB_(C^*β*^), where C^*α*^ and C^*β*^ are the contact maps of the two experimentally determined structures displayed in Fig. 1. In section 3.2, we combine our physics-based model with each term separately to create two hybrid models in which the SBM has a single basin of attraction. In section 3.3, we combine these potentials into a hybrid model with a dual-basin SBM.

### 3.2 Single-basin SBM

Combining the SBM defined by Eq. 4 with our physics-based model results in a potential energy function of the form,

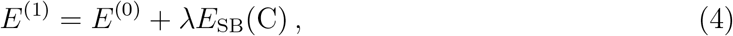

where *E*^(0)^ is the energy function defined in Ref [50] and λ is the strength of the structure-based term, which should be seen as a free parameter in this approach.

We apply Eq. 4 to the CTD of RfaH, with either *C* = C^*α*^ or C^*β*^ (see Fig. 1). Quite generally, we expect that increasing the relative strength of the SBM should increase the “nativeness” of the generated conformational ensemble. In other words, increasing λ should increase *Q_α_* in the case *C* = C^*α*^ and increase *Q_β_* in the case *C* = C^*β*^, where *Q_α_* and *Q_β_* are the fraction of native contacts formed with regards to the *α*-helical hairpin and *β*-barrel folds, respectively (see Methods). We indeed observe such a trend, as shown in Fig. 2. However, beyond this general trend, we find stark qualitative differences. The term *E*_SB_(C^*β*^) has a relatively large effect on the population of the all-*β* native state. For large enough λ, the melting curves are characterized by a sharp decrease in *Q_β_* with increasing temperature. Moreover, the folding transition occurs at increasingly higher temperatures, as indicating by a shift in the peaks of the heat capacity, *C*_v_. Interestingly, the height of the *C*_v_ peaks 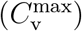 also increases with λ, suggesting also an increasing cooperativity of the transitions. By contrast, the single-basin SBM applied to the *α*-helical hairpin fold (*C* = C^*α*^) has a very different effect. The fraction of native contacts, *Q_α_*, does not reach 0.5, even at much lower temperatures. Strikingly, 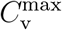 decreases with increasing λ in this case.

**Figure 2:**
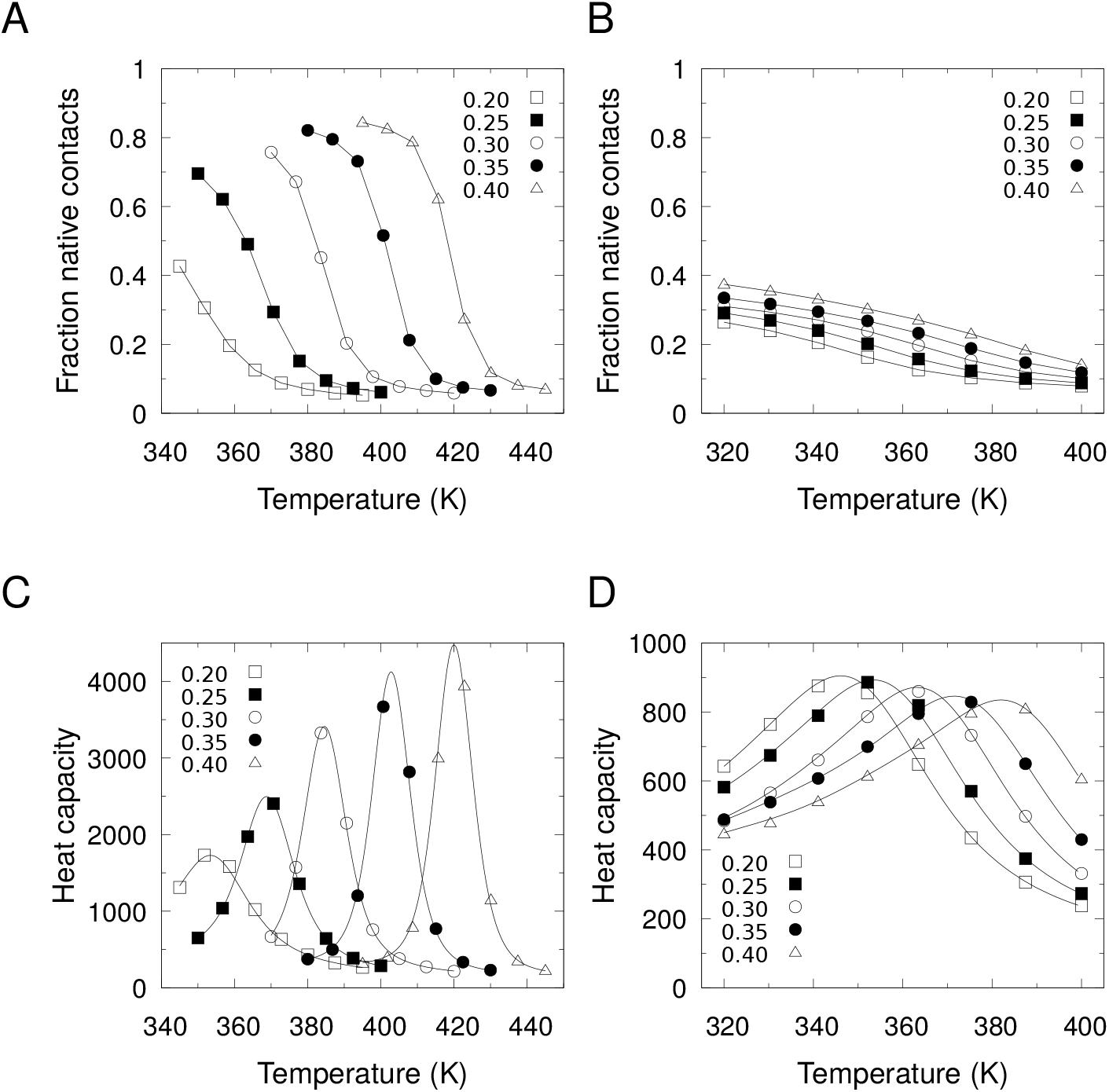
Equilibrium behavior of hybrid models with single-basin SBM. Shown are results obtained with the energy function in Eq. 4 with (A, C) a *β*-barrel SBM, *C* = C^*β*^, and (B, D) a *α*-helical hairpin SBM, *C* = C^*α*^, and the strengths λ = 0.20–0.40. As functions of the temperature: (A) *Q_β_*, (B) *Q_α_* and (C, D) heat capacity *C*_v_/*k*_B_, where *C*_v_ = (〈*E*^2^〉 – 〈*E*〉^2^)/*k*_B_*T*^2^, E is the energy from Eq. 4, and *k*_B_ is Boltzmann’s constant. Lines between points in (A) and (B) are drawn to guide the eye. Solid curves in (C) and (D) are obtained using multiple-histogram reweighting [58].

It must be noted that *E*^(1)^ in Eq. 4 does not preserve the energy scale of the underlying model [50], because *E*^(1)^ depends on λ. As a result, temperatures and energy quantities can therefore no longer be directly interpreted as real physical units. For example, obtained folding temperatures, *T*_f_, which we define using the *C*_v_ peaks, can in general not be compared with experimental data. However, comparing the Tfs for different proteins should still inform on their relative stabilities, as long as results are obtained for the same choice of λ.

### 3.3 Dual-basin SBM

We now combine the two structure-based terms, *E*_SB_(C^*α*^) and *E*_SB_(C^*β*^), into a dual-basin SBM. Following our general approach, the energy function of the resulting hybrid model becomes

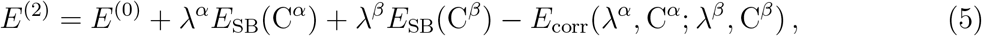

where strengths of the two structure-based terms, λ^*α*^ and λ^*β*^, are selected to be λ^*α*^ = λ^*β*^ = 0.30 based on the results of the previous section. The last term, *E*_corr_, is a correction term that is necessary in order to avoid a double counting of common contacts in C_*α*_ and C_*β*_ (see Fig. 1), which receive contributions from both *E*_SB_(C^*α*^) and *E*_SB_(C^*β*^). The role of *E*_corr_ is to eliminate the weaker of these two interactions (see Methods).

We apply Eq. 5 to the CTD as an isolated fragment and to the full-length RfaH. For the isolated CTD, we find that *Q_β_* increases with decreasing temperature, as shown in Fig. 3. In other words, despite the dual-basin nature of the SBM in our hybrid model, the isolated CTD folds into a stable *β*-barrel at low *T*. As a comparison, we show also in Fig. 3A the results for the single-basin SBM, which, interestingly, gives a higher stability compared to the dual-basin SBM. The simulations of full-length RfaH are carried out in the following way. First, the contact map C^*α*^ in Eq. 5 is replaced with C^*α*_RfaH_^, which in addition to all contacts in C^*α*^ also include NTD-CTD interdomain contacts, as shown in Fig. 3B. Second, for computational reasons, we keep the NTD backbone fixed in its native conformation while the linker and CTD regions are free to move. All sidechains are also left free. From these simulations we find, in contrast to the isolated CTD, that *Q_α_* increases with decreasing temperature. Hence, within the context of the full-length RfaH, the CTD switches into an *α*-helix rich state.

**Figure 3:**
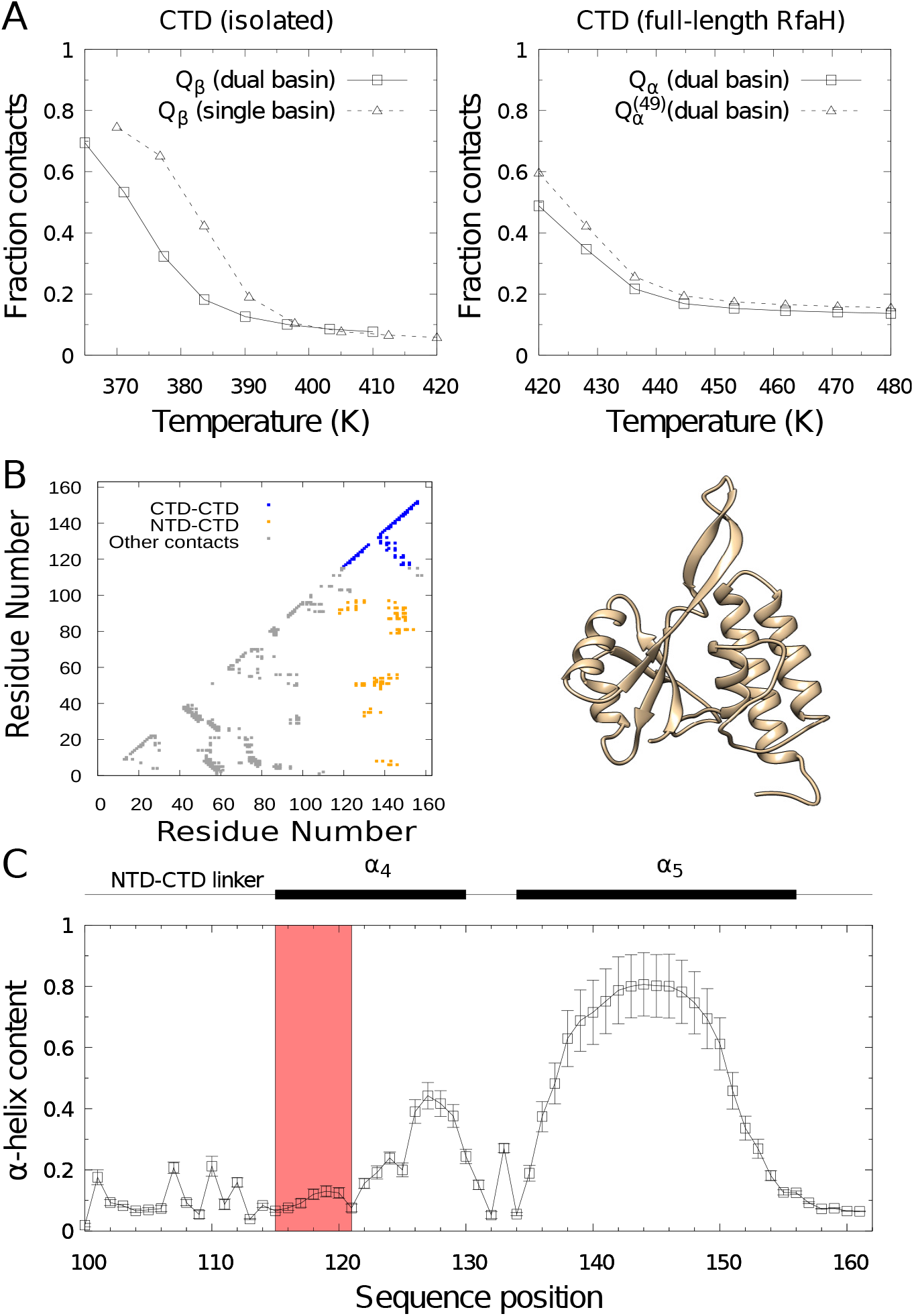
Equilibrium behavior of hybrid model with dual-basin SBM. (A) Temperature dependence of the fraction of native contacts (*Q_β_* or *Q_α_*) obtained from simulations of the isolated CTD or full-length RfaH. For comparison, results for the single-basin *β*-barrel SBM is re-drawn from Fig. 2A. as described below. (B) Structure in cartoon representation (right) and native contact map (left) of full-length RfaH (PDB ID 2oug). Intra-CTD, inter-domain and other native contacts are shown in blue, yellow and gray, respectively. Blue and yellow contacts make up the contact set, C^*α*_RfaH_^. (C) The average *α*-helix content as a function of sequence position for full-length RfaH, taken at 420 K (NTD region not shown). 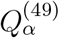 is determined as *Q_α_* but taken over a reduced set of 49 contacts that excludes contacts in the segment Val116-Gly121 (orange shade). All results are obtained with λ^*α*^ = λ^*β*^ = 0.30.

The fraction of native contacts, *Q_α_*, does not reach values higher than around 0.5 at the lowest simulated *T*, however, (see Fig. 3A). The examine the reason for this, we plot the propensity for forming *α*-helix structure as a function of residue position, as shown in Fig. 3C. The *α*_4_ helix (residues 116—130) is in our model significantly less structured than the helix (residues 134—156). According to the NMR experiments of Burmann et al. [21], the segment Val116-Gly121 displays chemical shifts that are more in line with a random coil than an *α*-helix, suggesting that the 6 N-terminal residues of *α*_4_ might be mainly disordered in the solution state. Therefore, we construct an alternative nativeness measure, 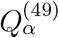, which is the same as *Q_α_* but all contacts involving 116-121 are ignored. Indeed, 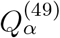 is consistently higher than *Q_α_* across all temperatures indicating that the *α*-helical hairpin fold is formed at low *T* with the exception of the N-terminal part of *α*_4_, which, in line with experiments, remain largely unstructured.

### 3.4 Folding energy landscape with a single dominant funnel

Having showed that our hybrid model with a dual-basin SBM captures the basic thermodynamic behavior of the RfaH CTD, i.e., its all-*α*-to-all-*β* switch in global free energy minimum upon excision from the rest of the protein, we turn to the folding behavior of this domain. To delineate the impact of the fold switching capability of the RfaH CTD, we compare with results from the hybrid model with a single-basin SBM (applied to the *β*-barrel contact map C^*β*^). Both these hybrid models fold the CTD into a stable *β*-barrel, but only the dual-basin SBM gives proper fold switching. We therefore reason that differences in the respective energy landscapes can be attributed to the unique fold switching behavior of the CTD.

We first determine the free energy surface *F*(*Q_α_, Q_β_*) close to the folding temperature, *T*_f_, as shown in Fig. 2A. At *T*_f_, one might expect our dual-basin SBM to produce three distinct free energy minima corresponding to the unfolded state, which must be substantially populated at *T*_f_, and two ordered states, the all-*α* fold and all-*β* folds. However, there are only two major minima in the *Q_α_-Q_β_* plane: (1) a low-*Q_α_*, low-*Q_β_* minimum, i.e., the unfolded state; and (2) a high-*Q_β_*, low-*Q_α_* minimum, i.e., the *β*-barrel state. The absence of a minimum for the *α*-helical hairpin fold is also clear from the the free energy surface *F*(*E*, RMSD_*β*_) in Fig. 4B, which show no distinct low-energy states that compete with the native *β*-barrel fold. At *T* < *T*_f_, i.e., under folding conditions, the population of the unfolded state is substantially decreased as ordered states with lower energies are increasingly favored. As a result, the energy landscape becomes dominated by a single funnel toward the *β*-barrel fold, as shown by the one-dimensional free energy profiles, *F*(*Q_α_*) and *F*(*Q_β_*), in Fig. 5.

**Figure 4:**
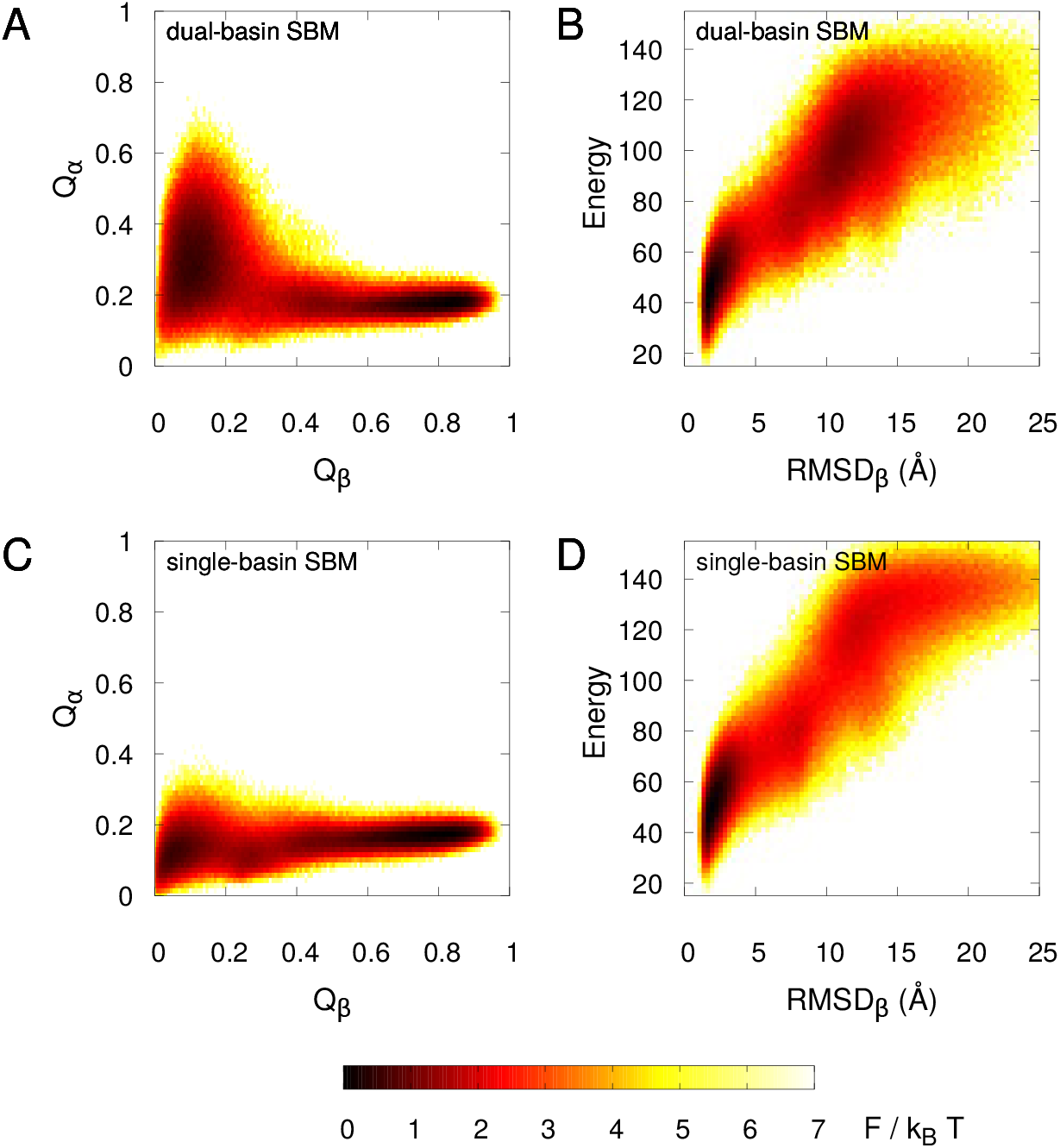
Free energy surfaces at the folding midpoint: single-basin SBM vs dual-basin SBM. Shown are free energy surfaces *F*(*X*_1_, *X*_2_) = - ln *P*(*X*_1_, *X*_2_), with (A, B) *X*_1_ = *Q_α_* and *X*_2_ = *Q_β_* or (B, D) *X*_1_ is the total energy and *X*_2_ = RMSD_*β*_. The probability distributions *P*(*X*_1_, *X*_2_) are obtained close to the respective *T*_f_s, which are (nominally) 384 K for the dual-basin SBM and 377 K for the (*β*-barrel) single-basin SBM.

**Figure 5:**
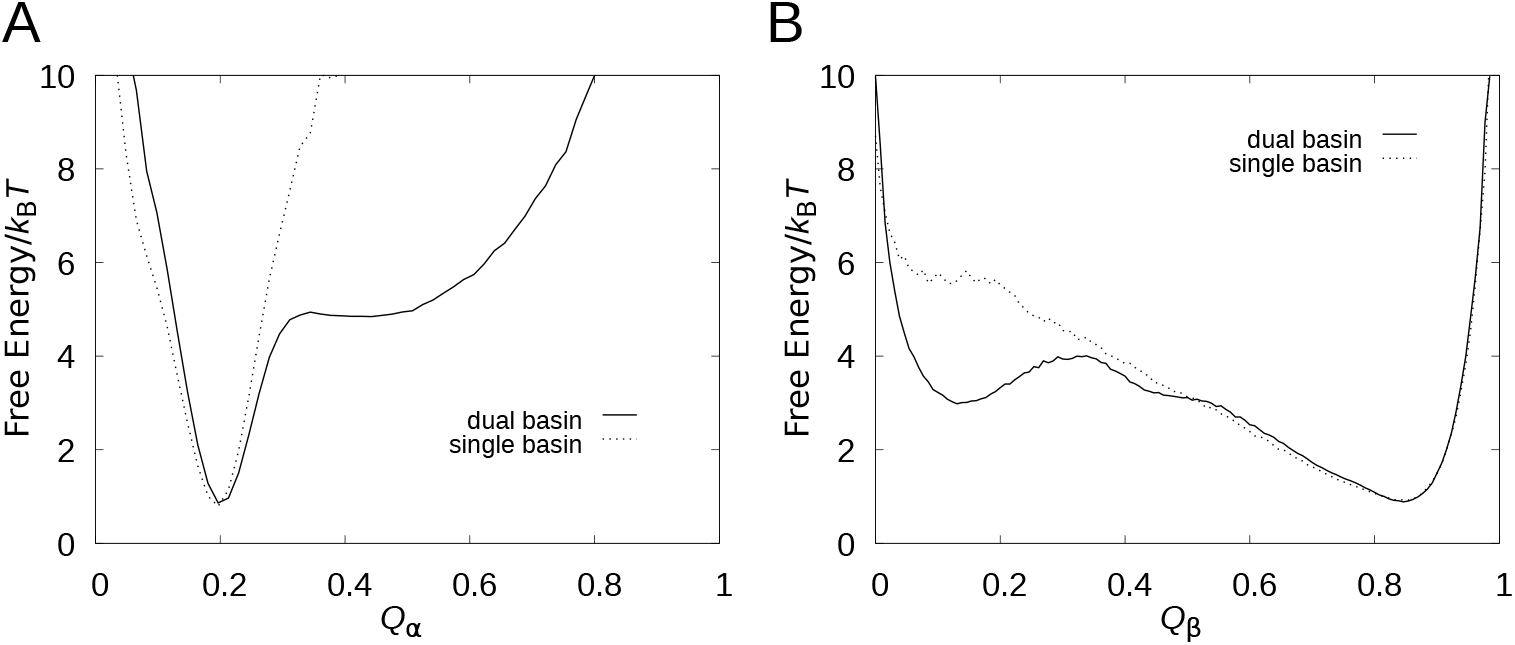
Free energy profiles under folding conditions. Shown is the free energy as function of (A) *Q_α_* and (B) *Q_β_* for hybrid models with either dual-basin (solid curves) or single-basin (dashed curves) SBM, taken at *T* ≈ 0.96*T*_f_.

Recent computational studies of the RfaH CTD have found energy landscapes with clear two-funnel character [32, 37], in apparent contradiction with our results. These studied relied on molecular dynamics simulations with either implicit or explicit water in combination with advanced sampling and analysis techniques. For example, Bernhardt et al. [37] achieved enhanced sampling using a Hamilton replica-exchange method that couples their physical model to a Gō potential, which seeds the sampling of conformational space. It is possible that these types of techniques are able to detect fine features of the energy landscape that are not apparent in our calculations. However, the lack of a major competing minimum corresponding to the *α*-helical hairpin fold in the energy landscape of the isolated CTD is consistent with the [^1^H,^15^N]-HSQC NMR data of Burmann et al. [21]. Upon protease-induced cleavage of the NTD-CTD linker, thus releasing the CTD into the solution, they observed a complete disappearance of the spectra from the *α*-helical hairpin and the emergence of the spectra from the *β*-barrel fold [21]. Although the *α*-helical hairpin is clearly encoded in the amino acid sequence of the CTD, these experiment results limit the possible “depths” of this funnel and hence its significance in comparison with the large number of other local minima present in any protein energy landscape.

### 3.5 The impact of encoding two folds on the unfolded state

While the differences between the free energy surfaces *F*(*Q_α_, Q_β_*) at *T* ≈ *T*_f_ (see Fig. 3A and B), obtained for the single- and dual-basin SBMs are not large, it is interesting to examine them in detail. We find that, for the single-basin SBM, the unfolded state is a rather narrow minimum, especially in the *Q_α_* direction. For the dual-basin SBM, the unfolded state is broader in the *Q_α_*-direction, indicating larger fluctuations. Many of the native contacts in C^*α*^ are local, such as *α*-helical i, i+4-contacts (see Fig. 1). The main impact of the encoding the *α*-helical hairpin within our dual-basin SBM, which nonetheless spontaneously fold into the *β*-barrel fold as we have shown, is to produce residual *α*-helix structure in the unfolded state. Indeed, we quantify the *α*-helix content in the unfolded state, as defined by *Q_β_* < 0.5, for both models and find that it is significantly higher for the dual-basin SBM (20%) than for the single-basin SBM (7.8%).

A similar picture is obtained from the free energy profiles *F*(*Q_α_*) and *F*(*Q_β_*) (see Fig. 5) at lower temperatures, *T* ≈ 0.96*T*_f_. We note, in particular, that the shape of *F*(*Q_β_*) is remarkably similar between the two models at large *Q_β_*. This similarity likely reflects that contacts unique to the *α*-helical fold are either highly unstable or otherwise impossible to form due to topological constraints, once folding has progressed towards the *β*-barrel beyond a certain point (roughly *Q_β_* ≈ 0.5).

### 3.6 Is there an activation barrier to the all-*α*-to-all-*β* fold switch?

Although we find in our hybrid approach, with a dual-basin SBM, a dominant funnel to the *β*-barrel fold, conformations sufficiently close to the *α*-helical hairpin will, by construction, be energetically biased towards this attractor. Therefore, a conformation prepared in this all-*α* state might still need to overcome an activation barrier before it can proceed downhill the energy landscape towards the *β*-barrel native state. Such a situation might occur *in vivo* when RfaH binds to RNA polymerase, which triggers the CTD to detach from the NTD [24].

In order to explore the potential barrier of fold switching in the all-*α*-to-all-*β* direction, we carry out small-step “kinetic” MC simulations. We consider two different starting points: (1) the *α*-helical hairpin and (2) an extended, open conformation. The temperature is held fixed at *T* = 365 Kelvin, for which the *β*-barrel is the thermodynamically dominant state (see Fig. 3). Hence, both starting points should eventually transform into the *β*-barrel fold, although the timescale of the transformation could be different. These two sets of simulations started from point (1) and (2) thus probe respectively the fold switching and folding of the RfaH CTD.

In Fig. 6, we show the relaxation of the *α*-helix content, 〈*α*〉, and the fraction of native contacts, 〈*Q_β_*〉, where 〈〉 indicates an average taken over 100 independent runs, towards their respective equilibrium values at this *T*, i.e., 〈*α*〉_equil_ = 0.08 and 〈*Q_β_*〉_equil_ = 0.70. In terms of 〈*Q_β_*〉, the folding simulations, which start at 〈*Q_β_*〉 = 0, rapidly “overtake” the fold switching simulations. This behavior indicates some degree of kinetic trapping early in the fold switching process. However, the associated barrier cannot be large because the two sets of simulations become statistical indistinguishable at around 10^6^ MC cycles, when 〈*Q_β_*〉 is still far from its equilibrium end point. A convergence of the two sets of simulations occurs also in terms of 〈*α*〉 at roughly the same point. Interestingly, and perhaps surprisingly, the folding simulations exhibit an initial increase in *α*-helix structure content to a maximum of 〈*α*〉 ≈ 0.25, and thereafter a much slower decrease following closely the trend of the fold switching simulations. This gradual decrease comes from the conversion of more and more of the trajectories into the *β*-barrel fold. Taken together, these simulations suggest a fold switching process from the *α*-helical hairpin state (N_*α*_) to the *β*-barrel state (N_*β*_) proceeding as N_*α*_ → U → N_*β*_, where U is the unfolded state of the CTD.

**Figure 6:**
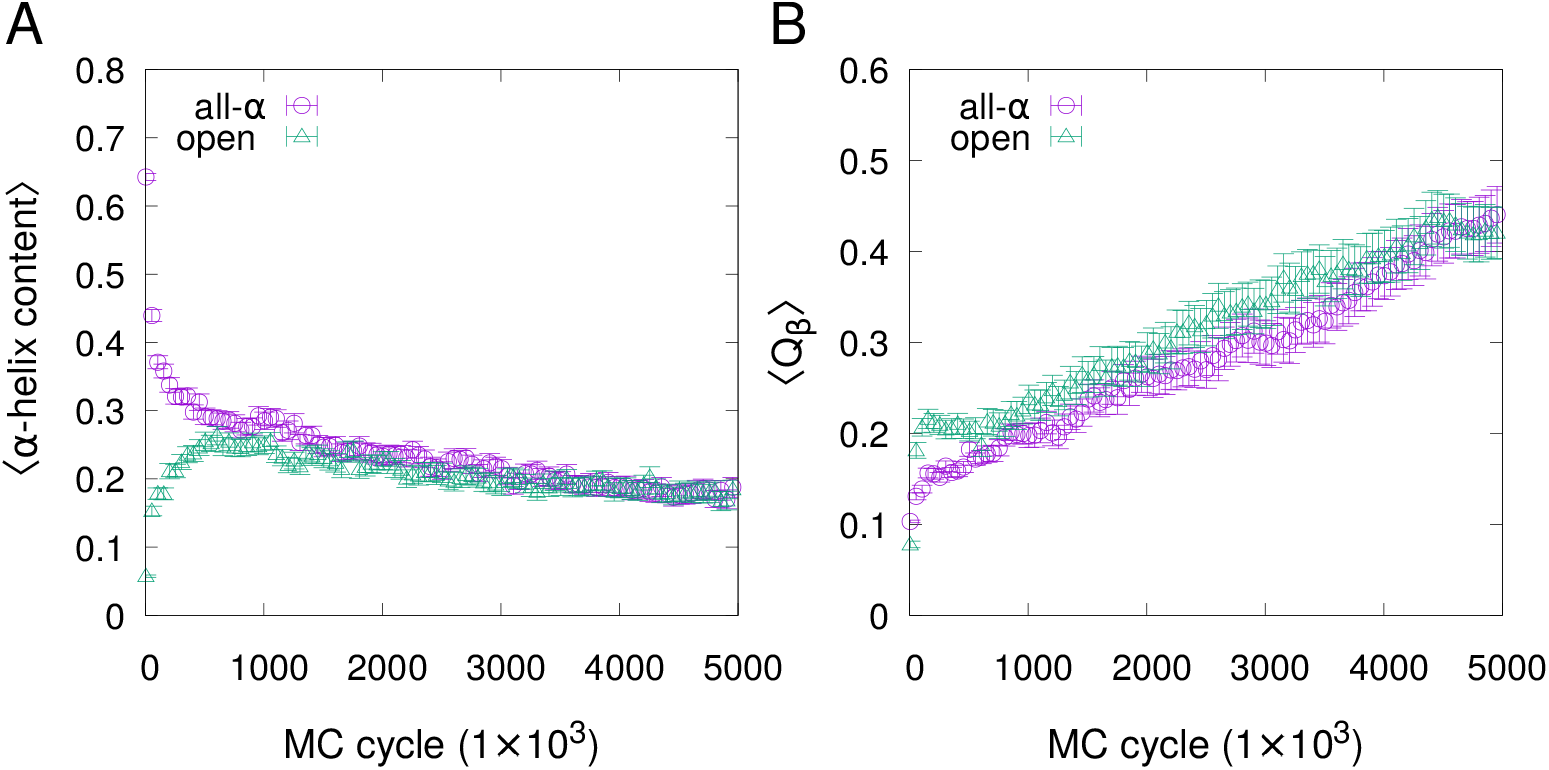
Fold switching and folding simulations. Shown is the evolution of the *α*-helix content, 〈*α*〉, and the fraction of native contacts, 〈*Q_β_*〉, in small-step Monte Carlo simulations started in either the *α*-helical hairpin fold (purple circles) or in an open chain conformation (green triangles), where the average 〈〉 is taken over 100 independent trajectories. Simulations are carried at *T* = 365 K. Error bars show standard errors.

This basic scheme is confirmed when examining in more structural detail individual fold switching events. Figure 7 shows RMSD_*α*_ and RMSD_*β*_ as functions of MC time for a typical fold switching trajectory, where RMSD_*α*_ and RMSD_*β*_ are the root-mean-square deviations taken with respect to the representative all-*α* and all-*β* CTD structures, respectively. After only a few MC cycles, RMSD_*α*_ increases rapidly, and the chain settles into an intermediate state with large conformational fluctuations. Residual *α*-helical structure are found upon inspection of conformations in this state, as shown in Fig. 7. Eventually, the chain switches abruptly to an all-*β* state, characterized by low RMSD_*β*_ values and much smaller chain fluctuations. Overall, we find that of all the trajectories that switch folds within the simulation time (47 out of 100), we find that none proceed directly from the all-*α* to the all-*β* state but instead proceed via an intermediate state. Because of the presence of residual *α*-helix structure, we identify this intermediate state with the unfolded state.

**Figure 7:**
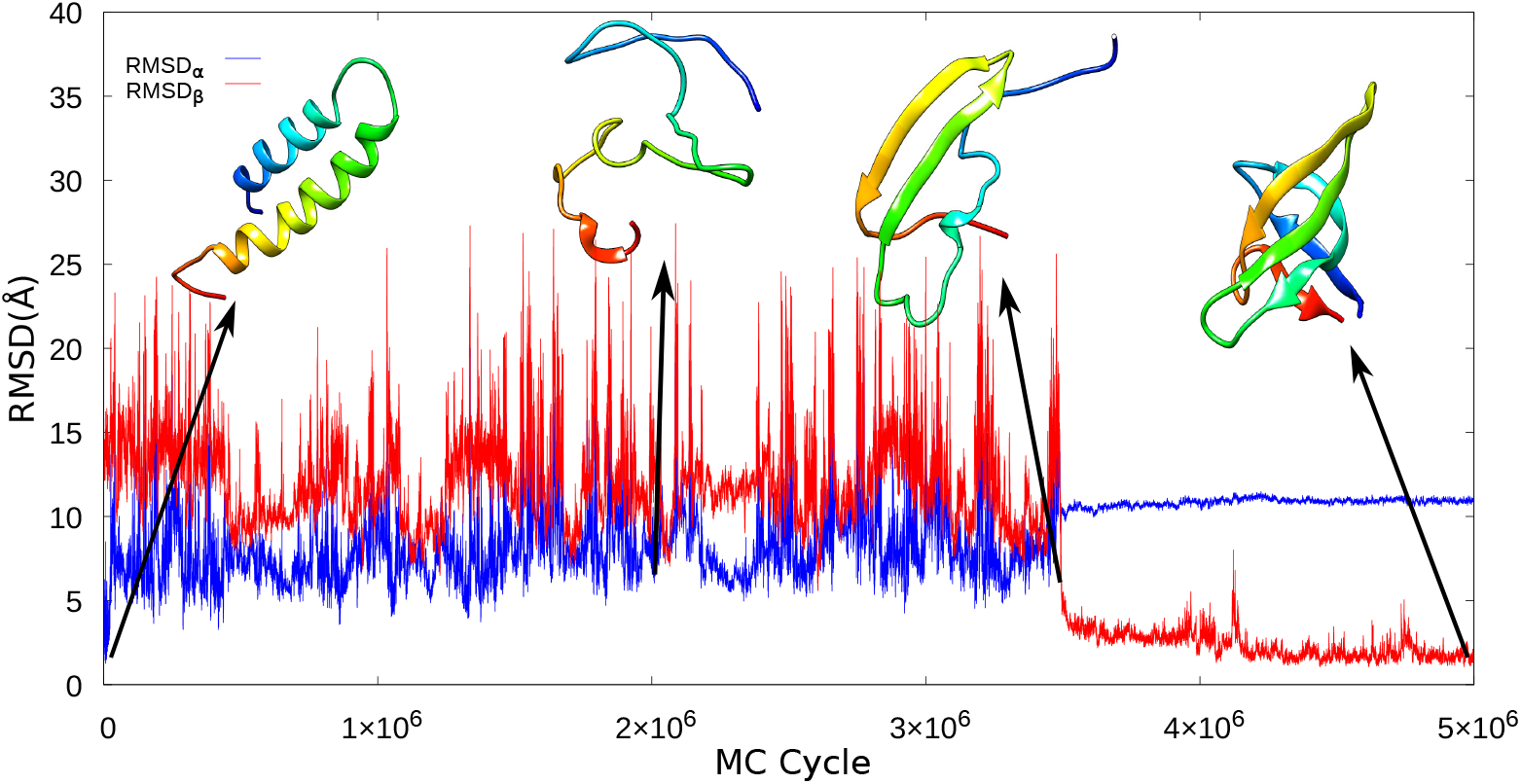
Example of fold switching trajectory. Shown is the evolution of the rootmean-square deviation determined with respect to the *α*-helix fold (RMSD_*α*_) or the *β*-barrel fold (RMSD_*β*_) for one of the 100 runs in Fig. 6.

## 4 Conclusions

Metamorphic proteins are often discussed in terms of an energy landscape with two separate and “coexisting” funnels in order to rationalize their ability to adopt two different folds. However, most naturally occurring metamorphic proteins adopt a unique fold under a given constant local environment and switch to a different fold only upon a change in the environment. To examine the energy landscape of the CTD of RfaH, we developed and tested a hybrid all-atom model that combines a physics-based model with a dual-basin structurebased potential (dual-basin SBM). We showed that this model captures the required change in global free energy minimum upon the excision of the CTD from RfaH. Applying this model to the isolated CTD, we found an energy landscape that is characterized by a single dominant funnel towards the *β*-barrel fold, with no sign of a second funnel towards the *α*-helical hairpin fold. Our model thus suggests that a multifunneled energy landscape cannot be assumed for metamorphic proteins. Further, by comparing with another hybrid model (with a single-basin SBM) that is similar but incapable of fold switching, we found that all major differences could be associated with the unfolded state. Specifically, this comparison indicated a relatively high content of *α*-helix structure in the unfolded state. Biophysical characterizations, e.g., using circular dichroism, of the CTDs of RfaH and other members of the general NusG/Spt5 family of transcription factors under weakly unfolding conditions, would provide an interesting experimental test of the computational results of this work.

## Acknowledgements

This work was supported by grants from Memorial University and Natural Sciences and Engineering Research Council of Canada (NSERC), and by computational resources provided by Compute Canada.

